# Predicting ovulation from brain connectivity: Dynamic causal modelling of the menstrual cycle

**DOI:** 10.1101/2020.08.11.247015

**Authors:** Esmeralda Hidalgo-Lopez, Peter Zeidman, TiAnni Harris, Adeel Razi, Belinda Pletzer

## Abstract

Longitudinal menstrual cycle research allows the assessment of sex hormones effects on brain organization in a natural framework. Here, we used spectral dynamic causal modelling (spDCM) in a triple network model consisting of the default mode, salience and executive central networks (DMN, SN, and ECN), in order to address the changes in effective connectivity across the menstrual cycle. Sixty healthy young women were scanned three times (menses, pre-ovulatory and luteal phase) and spDCM was estimated for a total of 174 scans. Group level analysis using Parametric empirical Bayes showed lateralized and anterior-posterior changes in connectivity patterns depending on the cycle phase and related to the endogenous hormonal milieu. Right before ovulation the left insula recruited the frontoparietal network, while the right middle frontal gyrus decreased its connectivity to the precuneus. In exchange, the precuneus engaged bilateral angular gyrus, decoupling the DMN into anterior/posterior parts. During the luteal phase, bilateral insula engaged to each other decreasing the connectivity to parietal ECN, which in turn engaged the posterior DMN. Remarkably, the specific cycle phase in which a woman was in could be predicted by the connections that showed the strongest changes. These findings further corroborate the plasticity of the female brain in response to acute hormone fluctuations and have important implications for understanding the neuroendocrine interactions underlying cognitive changes along the menstrual cycle.

## 1. Introduction

Natural fluctuations of ovarian sex hormones (i.e., estradiol and progesterone) affect the nervous system at multiple levels ^1^. Animal research has broadly evidenced the rapid changes exerted by sex hormones on the neuronal excitatory/inhibitory balance ^2^, synaptogenesis ^3^, myelination and re-myelination^4^. More importantly, these effects result in synaptic connectivity changes and therefore neural function ^5,6^, and in rodents a direct link to improved cognition has already been established for both hormones ^7,8^. However, in humans, menstrual cycle-related changes remain underinvestigated. Although sparse, acute structural changes related to hormonal fluctuations have already been reported for both grey ^9,10^ and white matter ^11^. Accordingly, connectivity patterns vary in women depending on the hormonal status, as shown by resting-state functional MRI (fMRI) ^12–14^. Amongst the most common approaches to resting-state functional MRI is the assessment of intrinsic connectivity networks (ICNs). These sets of brain networks with temporally correlated activity at rest relate to task-based BOLD activation patterns ^15,16^ and are consistent across healthy subjects ^17,18^. Three large-scale systems derived by independent component analyses (ICA) arise as ‘core’ neurocognitive networks, essential for cognitive functions ^19^. First, the default mode network (DMN), which includes the precuneus/posterior cingulate cortex (PCC), medial prefrontal cortex (mPFC) and bilateral angular gyrus (AG), is characterized by increased activity during the resting state and decreased activity during goal-directed tasks ^20,21^. Second, the salience network (SN), which comprises bilateral anterior insula (AI) and dorsal anterior cingulate cortex (ACC), is specialized in identifying and mapping relevant inputs, such as emotional stimuli ^22,23^. Third, the executive control network (ECN), is composed of bilateral middle frontal gyri (MFG) and bilateral supramarginal gyri (SMG) ^23^. This fronto-parietal system is usually lateralized ^16,17^ and responsible for higher cognitive control functions, such as working memory or directed attention, once the relevant stimuli are detected ^18,23^.

All of these ICNs are susceptible to modulation by sex hormone levels across the menstrual cycle. Within the DMN, intrinsic resting state connectivity has been shown to increase before ovulation ^9,24^ and decrease during the luteal phase to both left and right AG ^14,25^. Within the SN, higher activity and connectivity have been consistently reported during the luteal cycle phase and related to higher progesterone levels, both at rest and during tasks ^26,27^. Finally, for the ECN, increased task-based activation with higher levels of estradiol and progesterone have been shown for frontal areas. Specifically, the right MFG increased its activation during the luteal phase irrespective of the task^28–30^ (Hidalgo-Lopez et al., *under review*). Relatedly, during resting state, increased eigenvector centrality in bilateral dorsolateral PFC has been found in the presence of higher progesterone levels ^31^. Regarding inter-network connectivity, previous findings mainly report changes in the DMN-SN connectivity, pointing out an increase during the luteal phase (however, not without inconsistencies, see^24^). Specifically, ACC and amygdala (nodes of the SN) increased their connectivity with the precuneus (posterior DMN) during the luteal phase compared to menses ^32^. Likewise, when treated with progesterone, increased connectivity between several nodes of the SN-DMN has been reported ^26^. This enhanced inter-connectivity between SN and DMN, alongside the increased intrinsic connectivity in the SN and decreased intrinsic connectivity in the DMN, sets a very unique scenario for the luteal phase. Considering that both networks involved (specially ACC and mPFC) are also implicated in self-referential ^33^ and affective experience ^21,34^, this distinctive connectivity pattern could elicit the misattribution of salient stimuli and dysfunctional appraisal, leading to anxiety and depressive symptoms ^19^. Consequently, the coupling dynamics between and within the SN and DMN have been proposed to underlie the window of vulnerability for cycle-related affective disorders during the luteal phase ^27,35^. More importantly, the role of the ECN, responsible for the regulation of the SN and the DMN, still remains unexplored. Weis et al.^36^ observed decreased connectivity during the pre-ovulatory phase between the DMN and the left MFG (part of the ECN), while Petersen et al.^25^ observed decreased connectivity during the luteal phase between the ECN and the ACC (part of the SN), both compared to menses. These changes could imply a difference in functional integration of cognitive and affective processes, but up to now a cohesive model for understanding the global mechanisms in which the healthy female brain adapts to hormonal changes remains elusive. Despite being a useful resource to address brain organization, the assessment of ICNs derived from ICA ^16,18,21,37^ does not allow to infer the directionality of coupling between these multiple distributed systems. Effective connectivity methods, on the other hand, constitute the best approach to investigate the complex within and between ICN relationships across the different menstrual cycle phases. Specifically, model inversion with spectral dynamic causal modelling (spDCM) estimates hidden neural states from the observed Blood Oxygen Level Dependent (BOLD) signal – specifically, the cross spectral density of BOLD signals from different brain regions. It assumes that spontaneous fluctuations in the signal during resting state reflect the endogenous neural activity ^38^. By parameterising the hidden coupling among the neuronal populations, one can generate (complex) cross spectra among observed responses ^39–41^. In addition, spDCM is especially efficient to invert large DCMs of resting state fMRI ^41^, which makes it feasible to assess large-scale between and within networks’ organization.

In this paper, we aim to delineate the menstrual cycle-related changes in effective connectivity within and between DMN, SN and ECN, and identify the specific direction of previously reported effects. For example, it is yet not defined whether the enhanced inter-connectivity of the SN and DMN during the luteal cycle phase originates from the SN or the DMN. Likewise, it is unclear whether the downregulation of the SN during the luteal phase is related to an increase in the directed connectivity from the ECN as proposed by the triple network model. In addition, it also remains unexplored how the increased activation of the right MFG relates to increased afferent connectivity, given that the BOLD-response rather reflects the input to a neuronal population ^42^. As several psychiatric and neurological disorders share an aberrant intrinsic organization of the aforementioned three “core” networks ^19^, we consider it of the utmost importance to characterize their non-pathological directed organization in healthy women. In order to do so, we applied state-of-the-art effective connectivity analyses to a previously published longitudinal resting state-fMRI data set acquired during menses, pre-ovulatory and luteal cycle phase in a large sample of healthy young women ^14^. Finally, we aimed to identify, whether changes in specific directed connections were able to predict which cycle phase a woman was in.

## 2. Material and methods

### 2.1. Participants

A final sample of 60 healthy right-handed women aged 18-35 remained after exclusionary criteria. All of them had a regular menstrual cycle and did not use any hormonal contraceptives within the previous six months. Detailed demographics are provided in Hidalgo-Lopez et al.^14^. All participants gave their informed written consent to participate in the study. All methods conform to the Code of Ethics by the World Medical Association (Declaration of Helsinki) and were approved by the University of Salzburg’s ethics committee.

### 2.2. Procedure

Cycle duration was calculated based on participants’ self-reports of the dates of onset of their last three periods, and three appointments were scheduled once during menses (1–7 days after the onset of current menses; low progesterone and estradiol); once in the pre-ovulatory phase (2–3 days before the expected date of ovulation; peak estradiol, low progesterone), and once during the mid-luteal phase (3 days after ovulation to 3 days before the expected onset of next menses; high progesterone and estradiol), order counter-balanced. Pre-ovulatory sessions were confirmed by commercially available urinary ovulation tests (Pregnafix®). Participants had to confirm the onset of next menses in retrospect.

### 2.3. Hormone analysis

Saliva samples were collected from participants, stored and processed as described in Hidalgo-Lopez et al.^14^. Estradiol and progesterone were assessed using the Salimetrics High Sensitivity salivary Estradiol assay (sensitivity of 1 pg/ml) and the DeMediTec Progesterone free in saliva ELISAs (a sensitivity of 10 pg/ml), respectively. All samples were assessed in duplicates and samples with more than 25% variation between duplicates were reanalyzed.

### 2.4. Data acquisition

Functional images, fieldmaps and an MPRAGE sequence were acquired on a Siemens Magnetom TIM Trio 3T scanner. For the resting state we used a T2*-weighted gradient echo planar (EPI) sequence with 36 transversal slices oriented parallel to the AC–PC line (whole-brain coverage, TE = 30 ms, TR = 2250 ms, flip angle 70°, slice thickness 3.0 mm, matrix 192 × 192, FOV 192 mm, in-plane resolution 2.6 × 2.6 mm). Participants were instructed to close their eyes, relax and let their mind flow. For the structural images we acquired a T1-weighted 3D MPRAGE sequence of 5 min 58 sec (160 sagital slices, slice thickness = 1 mm, TE 291 ms, TR 2300 ms, TI delay 900 ms, FA 9°, FOV 256 × 256 mm).

### 2.5. Preprocessing

For the preprocessing, the first 6 images of each session were discarded, and functional images were despiked using 3d-despiking as implemented in AFNI (afni.nimh.nih.gov). The despiked images were then pre-processed using SPM12 standard procedures and templates SPM12 (www.fil.ion.ucl.ac.uk/spm) including segmentation of the structural images using CAT12. The resulting images were subjected to the ICA-AROMA algorithm implemented in FSL and non-aggressive removal of artefactual components ^43^.

### 2.6. Selection and extraction of volumes of interest

The ROIs were selected based on a large body of literature describing them as core nodes of the corresponding networks (Fig.1.a). Those included: precuneus/posterior cingulate cortex (PCC), bilateral angular gyri (AG) and medial prefrontal cortex (mPFC) for the DMN ^44,45^; bilateral anterior insula (AI) and anterior cingulate cortex (ACC) for the SN ^23,45^; and bilateral middle frontal gyri (MFG) and supramarginal gyri (SMG) for ECN ^17^. The ROIs and their group-level peak coordinates are listed in Table 1 and are also shown in Figure 1.a. Following Zhou et al., (2018), the group-level peaks were identified within each intrinsic connectivity network (ICN) using spatial ICA, as implemented in the Group ICA for fMRI Toolbox (GIFT, http://mialab.mrn.org/software/gift)^46^, and already described in Hidalgo-Lopez et al., (2020). After extracting 20 components, we selected those four corresponding to the posterior DMN (independent component; IC 19), posterior SN (IC 17), left ECN (IC 10) and right ECN (IC 11) via spatial correlation to pre-existing templates ^16^ (SI, Fig. 1). Both the DMN and the SN lacked of sufficient signal in anterior regions, and therefore, the anterior nodes of the DMN (mPFC) and SN (ACC) were identified through seed-based functional connectivity analysis from the posterior nodes (as done in Razi et al., 2015) using the CONN toolbox ^47^. The resulting areas were masked with BA10 & BA11 and ACC as implemented in the Wake Forest University (WFU) Pickatlas toolbox (Maldjian et al., 2003). Subject-specific coordinates were selected as local maximum within 8 mm of the group-level coordinates within the region-specific mask WFUpickatlas. For each of the 11 ROIs the principal eigenvariate from an 8 mm sphere around the subject-specific coordinates and within the ROI mask was extracted and corrected for CSF and WM (as done in Razi et al., 2015). These time series were then used in subsequent DCM analyses (Fig.1, b). Two participants were excluded due to insufficient signal in the mPFC.

**Table 1:** Group level volume of interest coordinates.

| Region | MNI coordinates |  |  | Network |
| --- | --- | --- | --- | --- |
|  | X | Y | Z |  |
| PCC | 3 | -55 | 31 | DMN |
| IAG | -45 | -64 | 31 | DMN |
| rAG | 46 | -63 | 34 | DMN |
| mPFC | 0 | 53 | -14 | DMN |
| IAI | -39 | 14 | -5 | SN |
| rAI | 40 | 11 | -5 | SN |
| ACC | 0 | 35 | 19 | SN |
| ISMG | -48 | -49 | 37 | ECN |
| rSMG | 51 | -46 | 37 | ECN |
| IMFG | -42 | 17 | 31 | ECN |
| rMFG | 42 | 20 | 34 | ECN |

**Abbreviation:** l, left; r, right; PCC: precuneus/posterior cingulate cortex; AG: angular gyrus; mPFC: medial prefrontal cortex; DMN: default mode network; AI: anterior insula; ACC: anterior cingulate cortex; SN: salience network; MFG: middle frontal gyrus; SMG: supramarginal gyrus; ECN: executive control network.

**Fig. 1.**
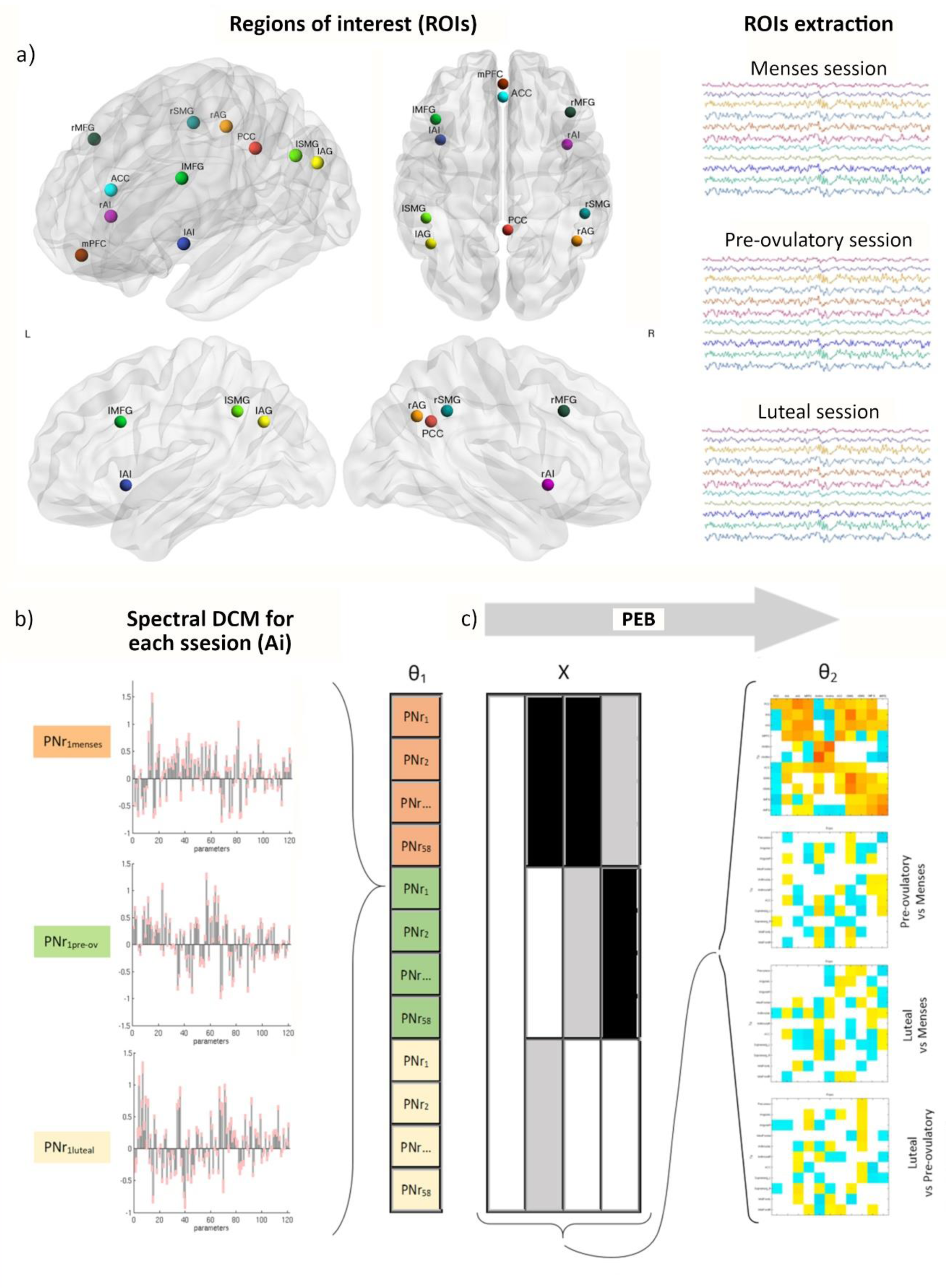
Procedures for dynamic effective connectivity analysis. The regions of interest from each ICN used in the current study is shown in (a). The default mode brain regions included the precuneus/posterior cingulate cortex (PCC), medial prefrontal cortex (mPFC) and bilateral angular gyrus (AG); the salience network comprised bilateral anterior insula (AI) and anterior cingulate cortex (ACC), and the executive control network was composed of bilateral middle frontal gyri (MFG) and bilateral supramarginal gyri (SMG). Each participant had three sessions locked to their menstrual cycle: during menses, pre-ovulatory and luteal. (b) A spectral DCM (spDCM) of 121 parameters was estimated for each session of every participant in a group DCM (θ_1_). (c) For the group level analysis, parametric empirical Bayesian analysis (PEB) was used to cycle phase group effects. This is a general linear model of the connectivity parameters. Shown is the design matrix X, where lighter colours indicate higher values. The estimated parameters are shown on the right. The columns are the outgoing connections, the rows are the incoming connections, ordered as: PCC, lAG, rAG, mPFC, lAI, rAI, ACC, lSMG, rSMG, lMFG, and rMFG. Hot colours indicate positive parameter estimates and cold colours negative.

### 2.7. Spectral Dynamic Causal Modelling and Parametric Empirical Bayes

Spectral DCM for resting state was specified and inverted using DCM12 as implemented in SPM12 (www.fil.ion.ucl.ac.uk/spm). For each participant and session, a fully connected model (including all possible connections between nodes), with no exogenous inputs, was specified to estimate the intrinsic effective connectivity (i.e., the ‘A-matrix’) within and between networks. The estimation fits the complex cross-spectral density taking into account the effects of neurovascular fluctuations as well as noise ^39^ and the default priors implemented in SPM were used at this level. The percentage of variance explained by the model for our subjects and sessions ranged from 91.60 to 99.27, which reflect good data fits for each model we estimated ^48^.

For the second level analyses, the parameters (effective connectivity strengths) were estimated in a Parametric Empirical Bayes (PEB) framework as described in Zeidman et al.^48^. To compute the difference in effective connectivity between the three different phases, three regressors were included: first, pre-ovulatory versus menses; second, luteal versus menses; and third, luteal versus pre-ovulatory (Fig.1, c). PEB results were thresholded to only include parameters from the A matrix that had a 75% posterior probability of being present vs absent, which represents a positive evidence for menstrual cycle-related changes (SI, Fig.2). Only results surviving this threshold are reported in the results section. We further assessed the hormonal modulation of the connections with a separate PEB analysis including scaled estradiol and progesterone levels, and their interaction as regressors. Results were thresholded to only include parameters that had a 75% posterior probability (SI, Fig.3). For brevity, they are not detailed in the results section, but summarized in SI Table 1.

### 2.9. Cross-validation

Finally, we assessed whether the individual cycle phase could be predicted based on the modulation of effective connectivity between those areas that survived a threshold of 99% posterior probability (very strong evidence) in the previous analyses. In order to do so we used a leave-one-out scheme (spm_dcm_loo.m) as described in Friston et al.^40^ and tested whether the effect on these particular connections could predict the cycle phase of participants.

## 3. Results

Results are displayed in Figures 2 and 3 and reported in SI Table 1.

**Fig. 2.**
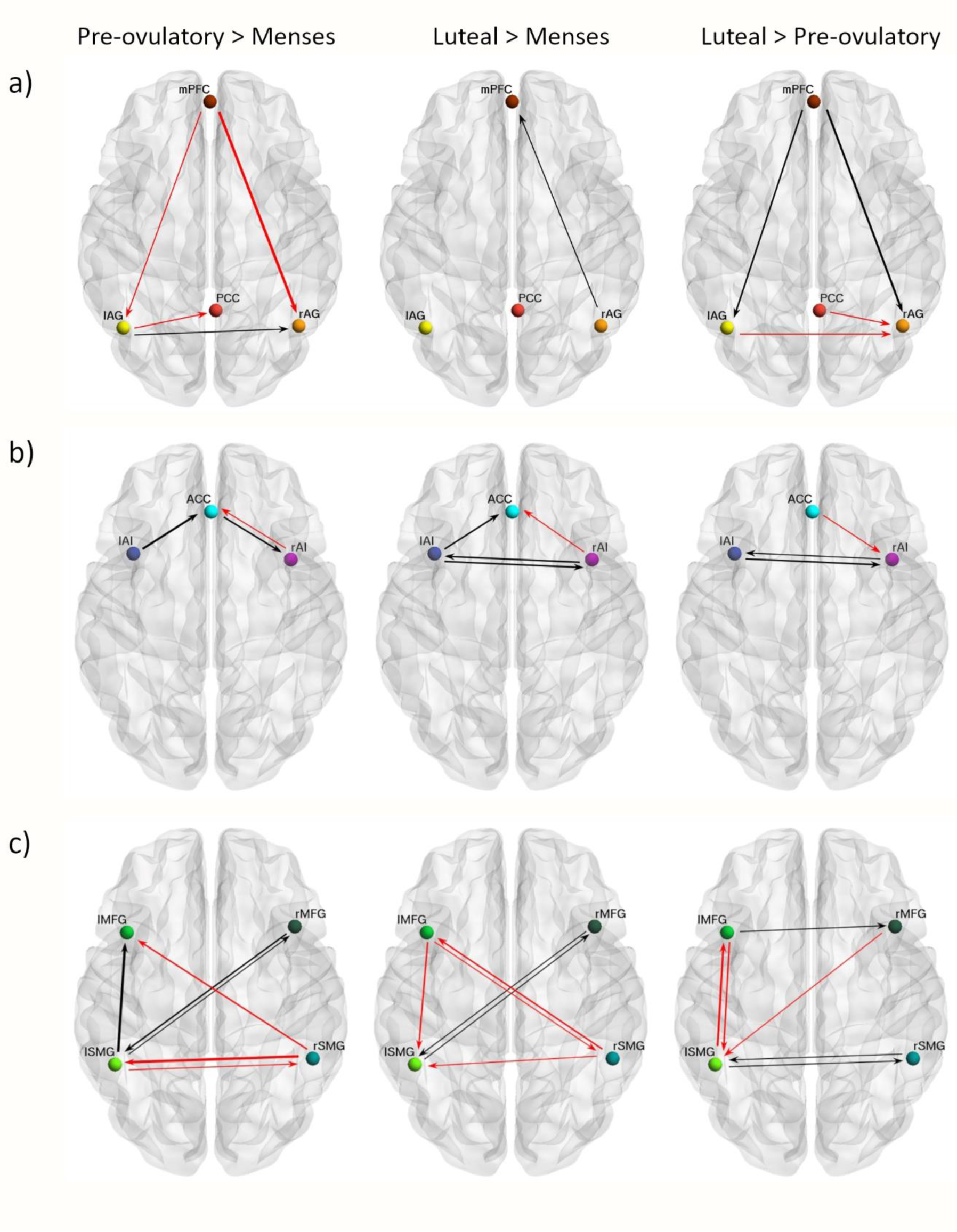
Cycle phase differences in within-network effective connectivity. A) DMN; B) SN; C) ECN. Only connections with a posterior probability > 0 .75 are displayed. The results reflect connection strengths as a difference between the indicated cycle phases. The differential connection strengths are depicted by the width of the arrow. Black arrows reflect positive values and red arrows reflect negative values for those connections which showed differences in the former than in the latter indicated cycle phase.

**Fig. 3.**
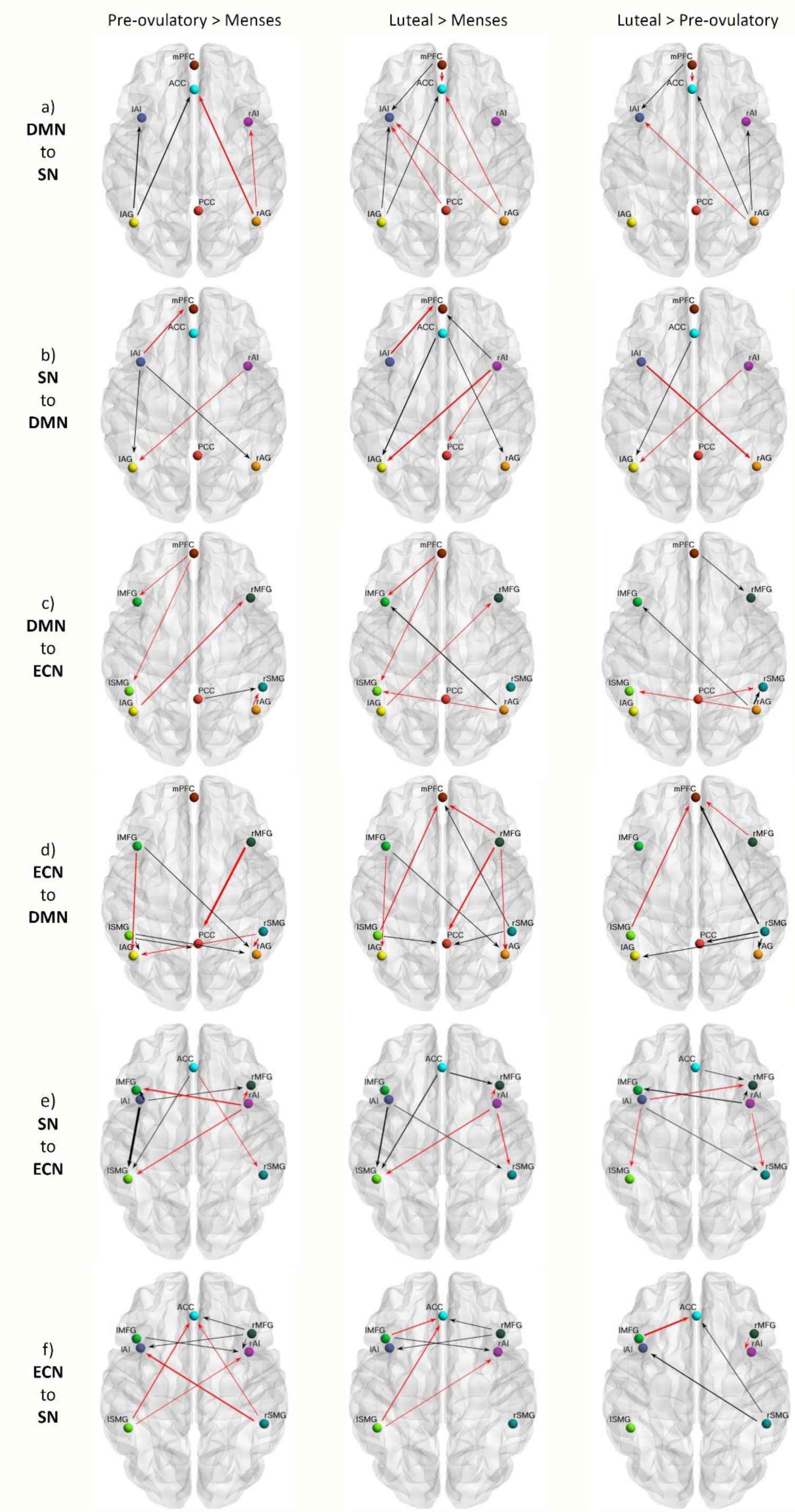
Cycle phase differences in between-network effective connectivity. A) DMN-SN; B) DMN-ECN; C) SN-ECN. Only connections with a posterior probability > 0.75 are displayed. The results reflect connection strengths as a difference between the indicated cycle phases. The differential connection strengths are depicted by the width of the arrow. Black arrows reflect positive values and red arrows reflect negative values for those connections which showed differences in the former than in the latter indicated cycle phase.

### Within-network effective connectivity

In general, effective connectivity within anterior and posterior nodes in the DMN was highest during menses and lowest during the pre-ovulatory phase (Fig.2.a). Within the SN bidirectional connectivity between left and right insula was strongest during the luteal phase. While effective connectivity from the left insula to the ACC was stronger during the pre-ovulatory and luteal phase compared to menses, effective connectivity from the right insula to the ACC followed the opposite pattern (Fig.2.b). Within the ECN, in general, connectivity between homotopic areas was weakest during the pre-ovulatory phase. Conversely, connectivity between the left lateralized nodes was the strongest right before ovulation (Fig.2.c).

### Between-network effective connectivity

Effective connectivity between the DMN and SN across the menstrual cycle was characterized by a lateralized pattern. From left AG to ipsilateral insula and ACC was stronger during the high-hormone phases, while from right AG to ipsilateral insula and ACC, connectivity was weakest during the pre-ovulatory phase. Furthermore, during the luteal phase, connectivity from the mPFC to left insula was the strongest, whereas to the ACC, the lowest (Fig.3.a). In general, effective connectivity from the SN to bilateral AG was strongest during the high-hormone phases, except for those connections originating in the right hemisphere. The effective connectivity from the SN to mPFC followed an opposite lateralization (Fig.3.b).

In general, effective connectivity from the DMN to the ECN was the lowest right before ovulation. During menses it was left lateralized, whereas during the luteal phase it was right lateralized (Fig.3.c). Effective connectivity from the posterior nodes of the ECN to the posterior DMN increased from menses to the luteal phase. During this phase, effective connectivity to the mPFC, from the left SMG was the lowest, whereas from the right SMG was the highest. During menses, connectivity was the strongest from frontal ECN to posterior DMN (Fig.3.d).

Effective connectivity changes from the SN to the ECN were also strongly lateralized. During the pre-ovulatory phase, connectivity from the left insula was the strongest, while from the right insula, in general was the lowest. Furthermore, effective connectivity from the SN to the right MFG was in general increased during the high-hormone phases (Fig.3.e). Effective connectivity from the frontal ECN to the SN was in general increased during the high-hormone phases. Effective connectivity from bilateral SMG to ACC and each contralateral insula decreased from menses to the pre-ovulatory phase. During the luteal phase, only those connections originating in the right hemisphere increased again (Fig.3.f).

### Prediction of cycle phase: cross-validation

We assessed whether individual cycle phase could be predicted based on the modulation of effective connectivity between those areas that survived a threshold of posterior probability > 0 .99 (very strong evidence) in the previous analyses. Those directed connections were from left anterior insula and right SMG to left SMG, from left SMG to left MFG, and from right MFG to PCC. The Pearson’s correlation coefficient was *r* = 0.21, *p =* 0.003 (Fig. 4). Thus, the difference across the menstrual cycle in effective connectivity between these areas was sufficiently large to predict the individual cycle phase. Effective connectivity from left insula to ipsilateral SMG, and from left SMG to ipsilateral MFG heavily increased from menses to the pre-ovulatory phase, and then decreased again during the luteal phase, but still staying higher than during menses. Conversely, effective connectivity from right MFG to PCC and from right to left SMG was stronger during menses than any other phase, decreased during the pre-ovulatory phase, and increased again during the luteal phase.

**Fig. 4.**
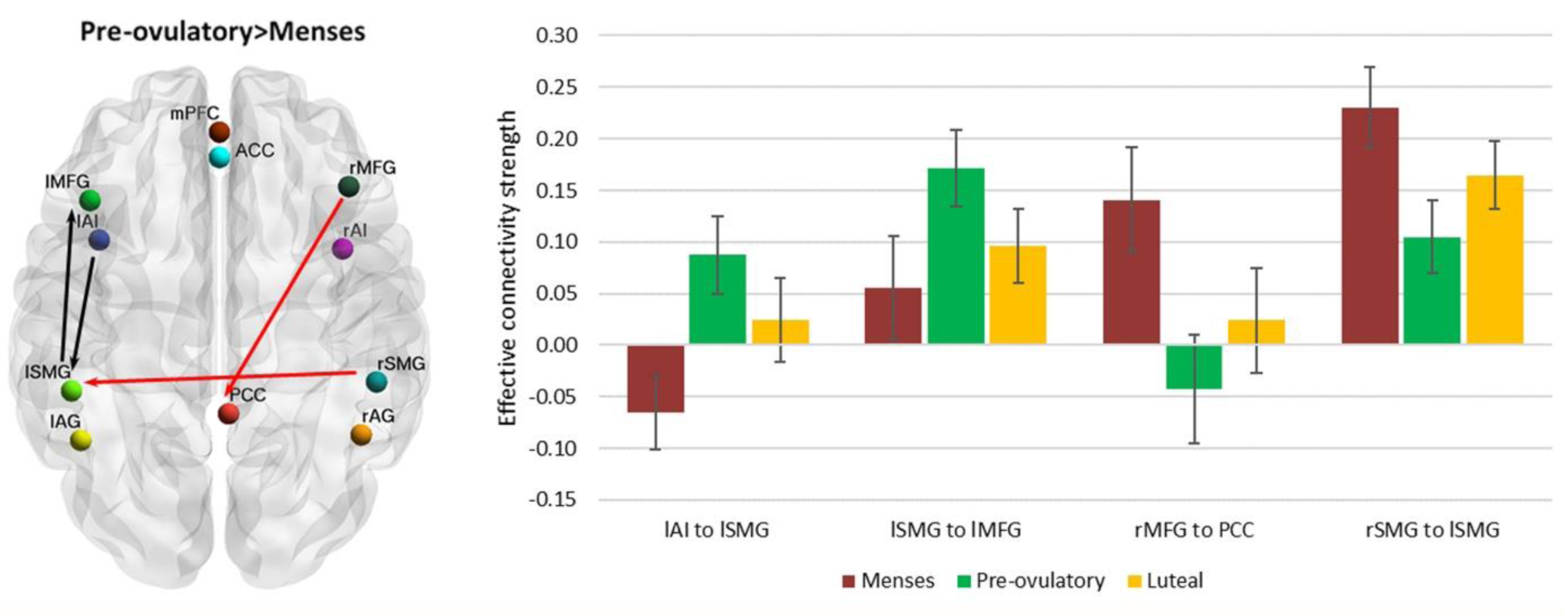
Cycle phase differences in effective connectivity within and between intrinsic connectivity networks DMN, SN and ECN with a posterior probability > 0 .99. These four connections predicted the cycle phase with a p<0.01. On the left, the differential connectivity strength between the indicated cycle phases is depicted by the width of the arrow. Black arrows reflect positive values and red arrows reflect negative values for those connections which showed differences in the former than in the latter indicated cycle phase. For displaying purposes, the individual values of each parameter were extracted and the mean is depicted for each cycle phase on the right bar chart. Error bars quantify the standard error of the mean.

## 4. Discussion

Research on the brain organization of naturally cycling women is of the utmost importance for understanding the neurobiological underpinnings of cognitive and emotional effects of sex hormones. However, to the best of our knowledge no prior studies have longitudinally assessed the resting-state effective connectivity related to the endogenous hormone fluctuations. Therefore, we used spectral DCM to characterize the temporal dynamics of brain connectivity in a triple network model across the natural menstrual cycle. Overall, some distinct patterns arose in each cycle phase and distinctively for each network, which will be discussed in detail in the following paragraphs and are depicted in Fig.5. In summary, during menses, which is characterized by low levels of estradiol and progesterone, we observed increased right lateralization of efferent connectivity from the SN and DMN, increased integration within the DMN and between DMN and ECN, and a higher recruitment of the SN by the parietal ECN. Then, right before ovulation, the lateralization shifted, as the left insula increased its efferent connectivity in response to heightened estradiol levels. It recruited the fronto-parietal network, which caused the right MFG to decouple from PCC. In exchange, the PCC engaged to bilateral AG, decoupling the DMN into anterior/posterior parts. Finally, during the luteal phase, the SN increased the connectivity within its own network, recruiting the right hemisphere again, and affecting differentially the other networks depending on the lateralization. Now the right insula recruited the frontal areas of the other networks. In turn, frontal nodes of ECN maintained the enhanced connectivity to the SN, while its posterior nodes increased their connectivity to the posterior DMN. In general, after ovulation, lateralization decreased as the homotopic regions of the ECN and SN were more connected to each other.

**Fig. 5.**
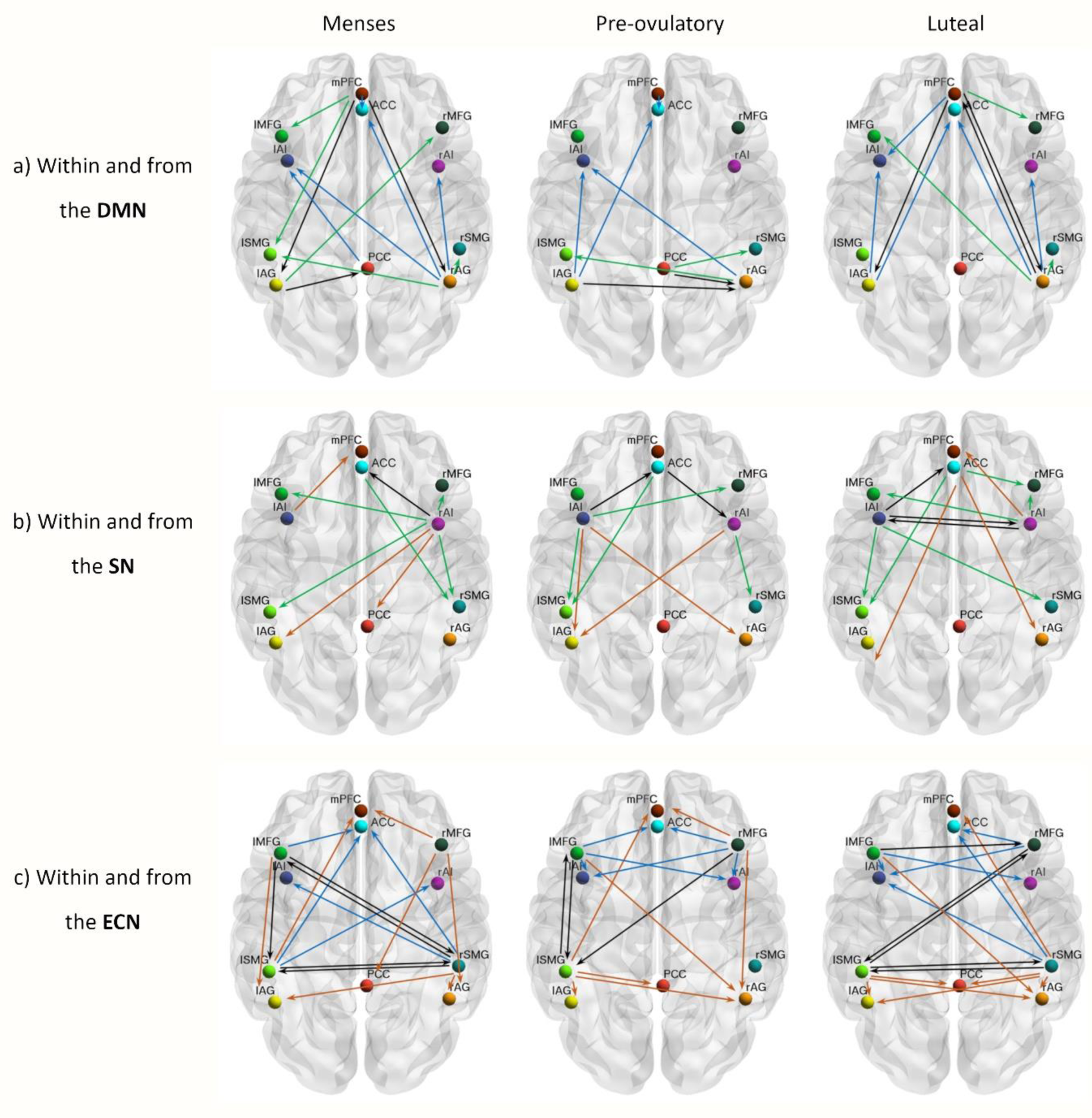
Summary of cycle-related differences in within and between-network effective connectivity. Each row depicts the connections observed to be enhanced in each cycle phase compared to the others within and from A) DMN; B) SN; and C) ECN. Only connections with a posterior probability > 0.75 are displayed. Within-network connections are depicted in black and the efferent connectivity to the DMN in brown, to the SN in blue, and to the ECN in green.

### DMN: endocrine anterior-posterior modulation

The present findings demonstrate a clear anterior-posterior modulation not only within the DMN but also in the connectivity between the DMN and the other networks. The global integration of the DMN was highest during menses, with enhanced connections from and within the entire DMN. During this phase, frontal ECN, particularly the left MFG, was most coupled with the DMN, in line with Weis et al.^36^. Right before ovulation, the bilateral AG disconnected from frontal areas and engaged to the PCC and to each other, increasing the connectivity within the posterior DMN. After ovulation, the AG switched again their connectivity to anterior areas disengaging from the PCC, and connections from the parietal ECN increased, especially to the posterior part of the DMN. The increased connectivity between bilateral AG and anteromedial areas during the luteal phase accompanied a decreased integrity of the posterior DMN, as the PCC decoupled from the DMN and was recruited by the parietal ECN. Remarkably, these findings suggest that the decomposition of DMN into anterior and posterior components, often described in ICA-analyses ^17,49^ at least in women, respond to hormonal factors.

The hypothesized increased connectivity between the DMN and SN ^27^ was only corroborated partially, given the anterior/posterior and lateralized pattern, and probably reflects an unbalanced mechanism rather than its organization in the non-pathological brain. Women with a history of affective side effects of the pill showed enhanced ACC-PCC connectivity both during treatment and the luteal phase compared to menses (not pill-active) ^32^. Furthermore, the over recruitment of the ACC into the DMN and enhanced connectivity with the mPFC has been characterized as unique for major depression symptoms and could reflect an inability to attend salient relevant external stimuli ^19^. Conversely, in our model we observed a reduced connectivity from mPFC to ACC during the luteal phase, related to higher progesterone levels (SI, table 1), while increased mPFC-PCC coupling to parietal areas, involved in external information processing. Thus, we suggest that these coupling-decoupling effects respond to a compensatory mechanism in healthy women, and its absence could underlie the vulnerability to depressive symptoms described during the luteal phase.

### SN: role of the insula as switcher in response to hormonal milieu

Increased reactivity of the SN, crucial for detecting emotional saliency, along with the deregulated coupling to frontoparietal systems has been proposed to underlie the vulnerability to develop affective disorders, in general ^19^, and particularly during the luteal phase ^27,35^. As expected, we found an increased within SN connectivity during the luteal phase related to higher hormone levels, reflected in the enhanced inter-hemispheric connectivity between the insulae (SI, table 1). Increased resting state functional connectivity between the insular cortices has already been related to higher estradiol levels in girls with precocious puberty ^50^. As a core node strongly and reciprocally connected to widespread cortical and subcortical areas ^51^, the insular cortex orchestrates interoception, cognition and emotion, contributing to emotional awareness, learning and memory processes ^51–53^. This unique position allows the anterior insula to engage to the ECN as it disengages from the DMN, acting as switcher between networks ^45,54,55^.

The present results not only corroborate previous findings in causal networks’ dynamics but also suggest a further aspect of the insula’s function related to its role in responding to changes in the endogenous hormonal milieu. Indeed, previous studies have already related the insula’s morphology and resting state connectivity to women’s hormonal status in a lateralized pattern ^56,57^. In our sample, the left insula had higher connectivity to the anterior DMN during menses, while the right insula was more strongly connected to the posterior DMN and ECN. Then, during the pre-ovulatory phase related to the increased estradiol levels, this pattern reversed and the left insula increased its connectivity to posterior DMN, ACC and ECN, as the right insula decreased it (SI, table 1). Although the bilateral anterior insulae are involved in the response to all emotional stimuli, a left-hemisphere dominance for positive stimuli ^58^ and empathy ^59^ has been suggested, relating the lateralized pattern of insula coupling and decoupling to the ECN to menstrual cycle dependent changes in emotion processing ^60^. During the luteal phase both insulae decoupled from the PCC as they increased the connectivity to each other. In addition, the left insula maintained the connectivity with parietal ECN while the right insula replaced its counterpart recruiting again the frontal ECN. Accordingly, these findings support the idea that the enhanced salience detection during the luteal phase could be buffered by coupling with frontal ECN. In fact, the increased connectivity between ACC and right MFG during the luteal phase may reflect an enhanced top down regulation in healthy women.

### ECN: lateralized and anterior-posterior hormonal modulation

Converging evidence from animal and human research relates estradiol to improved prefrontal-dependent functions, especially in verbal tasks (for a review, see ^5^). In women, frontal areas such as the IFG and MFG have shown enhanced activity during the pre-ovulatory phase and related to increased estradiol levels for implicit memory ^61^ and verbal encoding^62^. We recently found increased right frontrostriatal functional connectivity during resting state, related to higher estradiol levels during the pre-ovulatory phase ^14^. The increased afferent connectivity we observed from the left hemisphere to bilateral MFG right before ovulation could reflect underlying structural changes, but in humans, the relationship between molecular mechanisms and neural functions are difficult to determine. Nevertheless, evidence from animal studies strongly suggests the involvement of estrogen-dependent synaptogenesis in the PFC (in non-human primates^3^ and rodents^63^). Our model shows a strong shift of lateralization across the cycle phases with the most evident change being the recruitment of bilateral MFG from the left insula versus the right insula during menses and the luteal phase. In turn, the increased top-down engagement to SN from frontal ECN was maintained during the high-hormone phases. As expected, the right MFG increased its afferent connectivity during the luteal phase, including from left MFG, which could explain the increased executive-dependent activation previously observed (Hidalgo-Lopez & Pletzer, under review ^28–30^). Enhanced global network connectivity have been reported before in dorsolateral PFC, associated with increased levels of progesterone ^31^. Furthermore, overactivation of PFC ^64^ and differential insular activity ^65^ during high executive functions have been described for premenstrual dysphoric disorder. Specifically, in patients, the left insula was found less active than in healthy controls before ovulation, while more active during the luteal phase compared to both controls and pre-ovulatory patients. If, as our results suggest, the balanced bottom-up/top-down regulation entails a shift in the lateralization of the dorsolateral PFC afferent connections from the SN, its absence or deregulation could underlie some symptomatology.

After ovulation, parietal areas strongly coupled to the posterior DMN and to each other. Related to higher progesterone levels (SI, table 1), the right SMG increased the efferent connectivity to every node of the other networks but to right insula, which similarly decreased its connectivity to right SMG. A rightward asymmetry in SMG-insula connectivity has been found to be stronger in females than in males, and related to susceptibility to chronic pain disorders ^66^. In addition, the right SMG is particularly involved in shifting attention to salient stimuli ^67^. The shift in connectivity between the right insula and posterior ECN to anterior ECN after ovulation may reflect the dynamic integration of bottom-up/top-down processes, not only lateralized but also according to an anterior/posterior specialization.

### Lateralization

Across the menstrual cycle, two distinctive patterns regarding changes in lateralization could be distinguished. On the one hand, a right lateralized pattern during menses, which changes to the left hemisphere during pre-ovulatory, and recruiting the right hemisphere again during the luteal phase. This is the case for connectivity from SMG to posterior DMN, AG to SN, or the connectivity from the insula (broadly speaking). In addition, we observed an increased connectivity between homotopic regions of the SN and ECN during the luteal phase. Therefore, in most cases, this phase was characterized by a decreased asymmetry, which has been consistently reported and suggested to reflect a reduced transcallosal inhibition^68–70^. On the other hand, a more symmetrical pattern during menses and pre-ovulatory phase while right lateralized during the luteal phase is observed for the connectivity from ECN to the SN, in which the anterior-posterior modulation makes the scenario more complex. Furthermore, and as discussed previously, the right MFG also constitutes an exception to the lateralization observed across the menstrual cycle: its connectivity to the SN was the lowest during menses and increased as the hormone levels increased. These differential patterns could explain why the findings involving hemispheric asymmetries are sometime controversial and depend on the specific task and cognitive system involved ^71^.

### Prediction of cycle phase

Although the effect size of the cross-validation is small, the changes in connectivity strength of the four connections surviving a 99% threshold were sufficiently large to predict the individual cycle phase. Remarkably, all these connections reflect and summarize the effects that have been already discussed: increased engagement of the DMN with the ECN during menses, enhanced left frontoparietal connectivity right before ovulation, increased frontoparietal recruitment by the left insula in response to enhanced estradiol levels as well as interhemispheric decoupling in executive parietal areas, partially reversed after ovulation. Two common aspects should be noted. First, the most drastic changes occurred right before ovulation, which does not come as a surprise since the neuroendocrine feedback loops triggered by the surge of estradiol levels are quite unique in the human physiology, and the biological relevance of ovulation is undeniable. Structural changes between menses and the pre-ovulatory phase have already been shown to accurately classify cycle phase using a machine learning approach, and related to estradiol levels (BrainAGE^72^). Second, all four connections involved at least one node from the ECN, reflecting the key role of this network in menstrual cycle-related changes and SN/DMN coupling dynamics. A more detailed discussion of the functional role of each of these connections can be found in the supporting information, “Prediction of cycle phase” section.

### Final remarks

In conclusion, both lateralization and anterior-posterior effective connectivity patterns depend on the endogenous hormonal status in healthy young women. Remarkably, the specific cycle phase in which the woman was in could be predicted by some of the connections, which are representative of each unique phase pattern. This further corroborates the plasticity of the brain in response to the acute ovarian hormone fluctuations across the natural menstrual cycle. It also corroborates a differential effect of each hormone depending on the brain region and network dynamics and remarks the relevance of widening the focus to large-scale systems interaction, rather than the activity of localized brain areas. In fact, the cycle-related effects on connectivity from different nodes of a network to other networks suggest that ICA, the most used approach up to now, might be too coarse to capture the hormonal effects since the time-courses of various areas are averaged together and the results depend on the areas included in the ICN. Therefore, we propose a triple network dynamic model of menstrual cycle-related changes, with significant implications for understanding the underlying neuroendocrine interactions. This hormonal modulation may not only mediate menstrual cycle-related disorders, but also neurological and psychiatric changes across the menstrual cycle.

## 5. Acknowledgements

The authors want to thank the students of Belinda Pletzer for their assistance during participant’s recruitment and data acquisition, and the Pregnafix®company ATT Drogerievertriebs GmbH, for donating ovulation tests. We also thank all participants for their time and willingness to contribute to this study. Furthermore, we would like to thank the support of The Wellcome Centre for Human Neuroimaging, University College London.

## 8. Author Contributions

BP designed and made the concept of the study. EH and TH were responsible for data acquisition. Analysis of the data was performed by EH, and revised by PZ and AR. Interpretation of the results was done by EH and BP. EH drafted the manuscript, which was revised and approved by BP, PZ and AR. All authors agree to be accountable for all aspects of the work in ensuring that questions related to the accuracy or integrity of any part of the work are appropriately investigated and resolved.

