## SUPPORTING INFORMATION for "Predicting ovulation from brain connectivity: Dynamic causal modelling of the menstrual cycle"

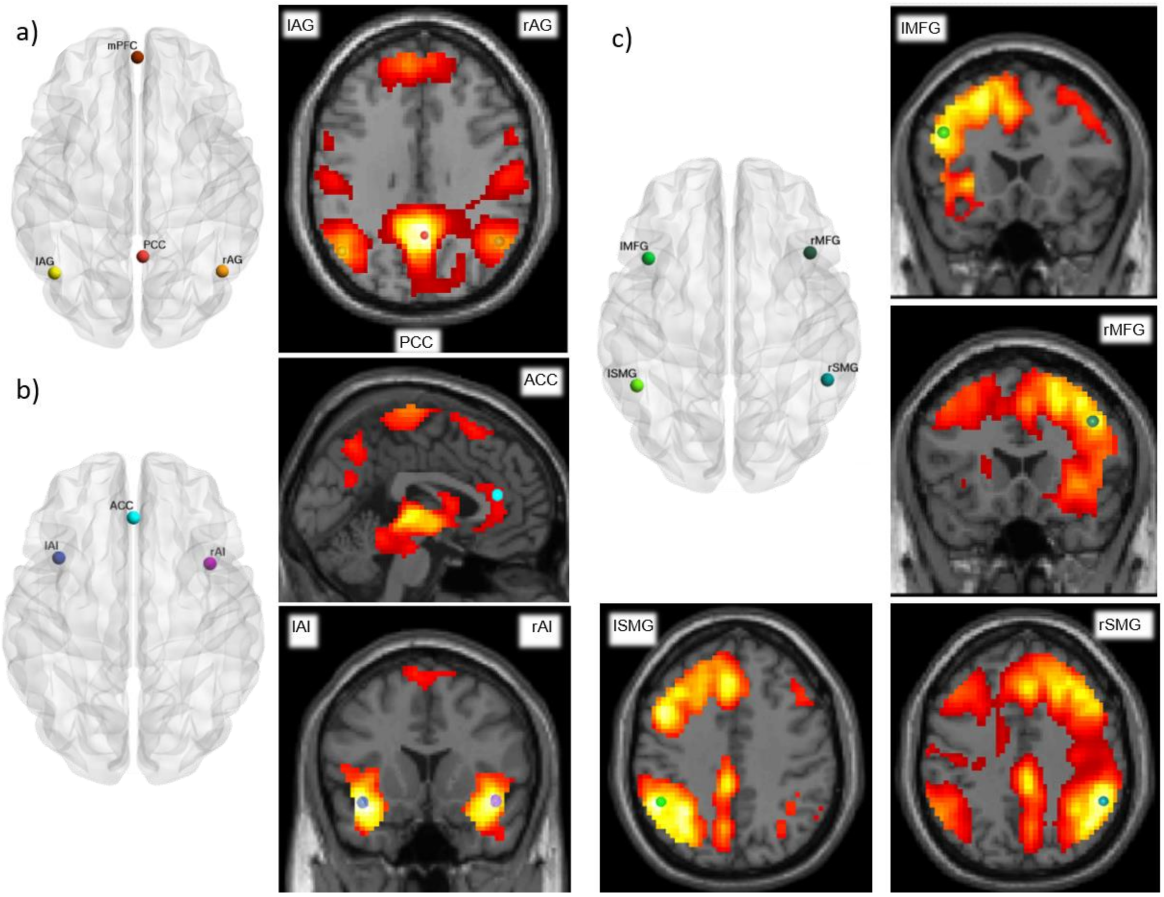


**Fig. 1:** Group-level ROIs identified using spatial independent component analysis (ICA). The group-level peak coordinates are overlaid on the spatial distribution maps from the ICNs of interest, correspondent to the DMN, SN, left ECN and right ECN, as spatially matched to pre-existing templates from Laird et al., 2011.

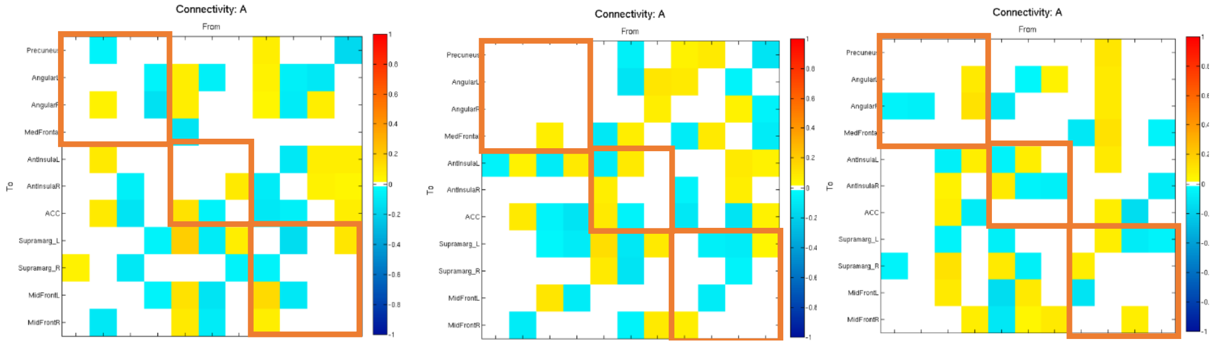


**Fig. 2:** Effective connectivity matrix reflecting differences in effective connectivity within and between intrinsic connectivity networks DMN, SN and ECN across the menstrual cycle. The columns are the outgoing connections, the rows are the incoming connections, ordered as: PCC, lAG, rAG, mPFC, lAI, rAI, ACC, lSMG, rSMG, lMFG, and rMFG. Hot colours indicate positive parameter estimates and cold colours negative. Only connections with a posterior probability > 0 .75 are displayed. A) Pre-ovulatory vs. menses, b) luteal vs. menses, c) luteal vs. pre-ovulatory.

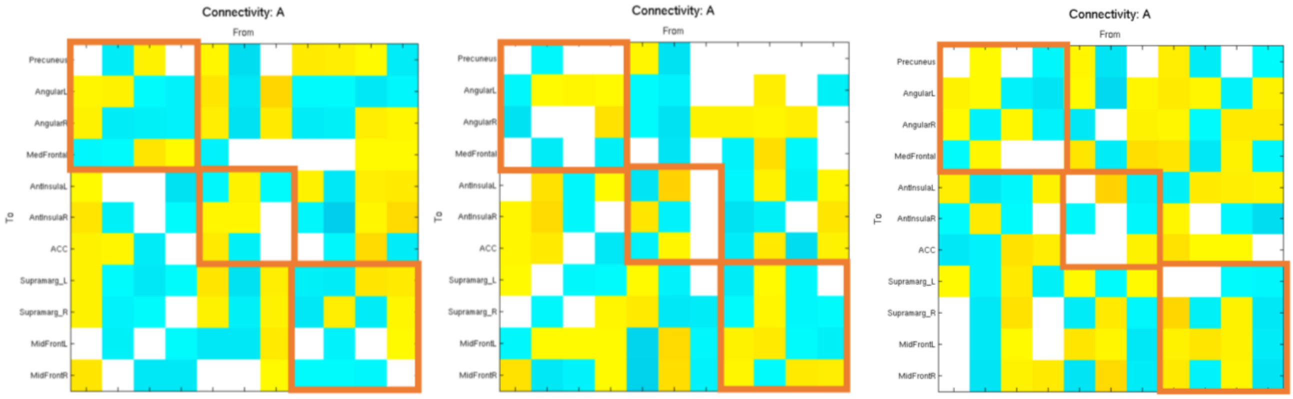


**Fig. 3:** Effective connectivity matrix reflecting differences in effective connectivity within and between intrinsic connectivity networks DMN, SN and ECN related to hormone levels a) estradiol, b) progesterone, and c) its interaction. The columns are the outgoing connections, the rows are the incoming connections, ordered as: PCC, lAG, rAG, mPFC, lAI, rAI, ACC, lSMG, rSMG, lMFG, and rMFG. Hot colours indicate positive parameter estimates and cold colours negative. Only connections with a posterior probability > 0 .75 are displayed.

***Detailed discussion of connections predicting menstrual cycle phase***

From left insula to the left SMG: The role of the insula identifying relevant internal and external stimuli and recruiting frontoparietal areas to guide behaviour would be essential right before ovulation for reproductive purposes. Relatedly, estradiol modulates temporal decision-making and impulsivity (Diekhof, 2015) which has been associated to the insula activity. As previously described, healthy women showed increased left insula activation during response inhibition right before ovulation compared to premenstrual dysphoric disorder patients (Bannbers et al., 2012). The strong increase of effective connectivity from the left insula to the left SMG, could explain changes in impulsivity that have been observed before ovulation for some women. In a previous work, we found interindividual differences in left putamen connectivity to ACC and left SMG right before ovulation, and related to changes in inhibitory control across the menstrual cycle (Hidalgo-Lopez and Pletzer, 2019). Given that striatal-insula functional connectivity is related to impulsivity trait (McHugh et al., 2013), the present model would place the left insula as intermediate between subcortical and cortical structures. The enhanced efferent connections of the left insula during the pre-ovulatory phase could explain the absence of interindividual differences in BOLD-activation, but opposite patterns of connectivity between left striatum, ACC and left SMG depending on the impulsiveness of women.

From left SMG to left MFG: In turn, the left SMG increased its connectivity to the left MFG. Both areas are structurally connected by the superior longitudinal fasciculus, previously reported to vary depending on sex hormones levels (Herting et al., 2012). Interestingly, both areas are the structural underpinnings of the human mirror system, and related to language and verbal abilities (Caspers et al., 2010; Farina et al., 2020). The mirror system plays an important role in social cognition and emotional empathy, and its sex differences have been suggested to depend on the hormonal milieu (Farina et al., 2020). Higher accuracy in emotion discrimination and recognition has been related to increased connectivity and activation within the mirror system (Farina et al., 2020). The increased input in left MFG from left parietal areas during the pre-ovulatory phase, caused in turn by the left insula, could underlie a general better emotion recognition described for this phase (Poromaa and

Gingnell, 2014). Likewise, given that the left MFG corresponds to the Broca's area, the increased connectivity could also underlie improved verbal abilities related to higher hormonal levels (for a review, see Luine, 2014; Sherwin, 2012).

From right MFG to PCC: On the other hand, the right MFG strongly disengaged from the PCC in response to the rise of estradiol levels, which allows the later to couple to posterior areas of DMN and ECN. Dorsolateral PFC and PCC are strongly interconnected and have been suggested to underlie (specially the right hemisphere) the necessary associative processes for self-awareness, self-representation and conscious experience (Cavanna and Trimble, 2006). The effective connectivity between these areas has already been reported to be modulate by endocrine factors in males, decreasing from dorsolateral PFC to PCC after oxytocin treatment (Kumar et al., 2019). In women, structural changes in the PCC related to the hormonal milieu have also been reported, with decreased cortical thickness in oral contraceptive users (Petersen et al., 2015).

From right SMG to left SMG: Finally, while the driving input for left SMG during menses is its homotopic region, the rise in estradiol levels causes an interhemispheric decoupling, allowing the left hemisphere to increase its within connectivity. The SMG is involved in cross-modal integration processes, comprising a wide range of functions such as proprioception, calculation, social cognition, or praxis functions. In the later, a left predominance for planning and verbal communication, while right one for visuospatial processing is well-known. For the execution of the action itself, the lateralization is more diffused (Bohlhalter et al., 2009). Menstrual cycle-related changes in the connectivity between homotopic SMG could underlie the differential effect of sex and hormonal status during visuo-spatial tasks. During menses, women showed more activation in the left SMG, and did not differ behaviourally from men (Schöning et al., 2007). After ovulation, progesterone counteracted estradiol, and the homotopic connectivity increased again, contributing to a decreased asymmetry.

| | From menses to pre-ovulatory | From menses to luteal | From pre-ovulatoy to luteal | E | P | E*P |
|---|---|---|---|---|---|---|
| *Negative impact of higher estradiol and/or progesterone levels and combinatory effect:* | | | | | | |
| From left AG to PCC | Decreased | | | - | - | + |
| From left AG to right AG | Increased | | Decreased | - | | - |
| From left AG to right MFG | Decreased | Decreased | | - | - | - |
| From right AG to left AI | | Decreased | Decreased | | - | - |
| From right AG to left SMG | | Decreased | Decreased | - | - | + |
| From mPFC to ACC | | Decreased | Decreased | | - | + |
| From mPFC to left SMG | Decreased | Decreased | | - | - | - |
| From left AI to left MFG | Increased | | Decreased | - | - | + |
| From right AI to PCC | | Decreased | | - | - | - |
| From right AI to right SMG | | Decreased | Decreased | - | - | + |
| From right AI to left SMG | Decreased | Decreased | | - | - | - |
| From right AI to left AG | Decreased | Decreased | Decreased | - | - | - |
| From left SMG to mPFC | | Decreased | Decreased | | - | + |
| From left MFG to left AG | Decreased | Decreased | | - | | - |
| From left MFG to right SMG | | Decreased | | - | - | + |
| **From right MFG to PCC** | **Decreased** | **Decreased** | | **-** | | **-** |
| *Negative impact of higher estradiol and/or progesterone levels is partially reversed by the combinatory effect:* | | | | | | |
| From right AG to ACC | Decreased | Decreased | Increased | - | | + |
| From right AG to right AI | Decreased | | Increased | | - | - |
| From right AG to right SMG | Decreased | | Increased | - | | + |
| From left SMG to right AI | Decreased | Decreased | | - | - | + |
| From left SMG to ACC | Decreased | Decreased | | | - | + |
| From left SMG to right SMG | Decreased | | Increased | - | - | + |
| *Negative impact of higher estradiol levels is reversed by progesterone positive impact and combinatory effect:* | | | | | | |
| From mPFC to left AG | Decreased | | Increased | - | + | - |
| From mPFC to right AG | Decreased | | Increased | - | + | - |

| | | | | | | |
|---|---|---|---|---|---|---|
| From mPFC to left MFG | Decreased | Decreased | | - | + | |
| From mPFC to right MFG | | | Increased | - | + | + |
| From left AI to mPFC | Decreased | Decreased | | - | | + |
| From right AI to ACC | Decreased | Decreased | | - | + | |
| From right AI to left MFG | Decreased | | Increased | - | + | + |
| From right AI to right MFG | Decreased | Decreased | Increased | | + | + |
| From right SMG to left AG | Decreased | | Increased | - | + | + |
| From right SMG to right AG | Decreased | | Increased | - | + | - |
| From right SMG to mPFC | | Increased | Increased | | + | - |
| From right SMG to left AI | Decreased | | Increased | - | + | + |
| From right SMG to ACC | Decreased | | Increased | - | + | + |
| **From right SMG to left SMG** | **Decreased** | **Decreased** | **Increased** | **-** | **+** | |
| From right SMG to left MFG | Decreased | Decreased | | - | + | + |
| From left MFG to right MFG | | | Increased | - | + | + |
| *Positive impact of higher estradiol and/or progesterone levels and combinatory effect:* | | | | | | |
| From right AG to left MFG | | Increased | Increased | | + | + |
| From right AG to mPFC | | Increased | | + | | |
| From left AI to right AI | | Increased | Increased | + | + | - |
| **From left AI to left SMG** | **Increased** | **Increased** | **Decreased** | **+** | **+** | **+** |
| From left AI to right SMG | | Increased | Increased | + | + | - |
| From right AI to left AI | | Increased | Increased | + | + | + |
| From ACC to left AG | | Increased | Increased | + | | + |
| From ACC to right AG | | Increased | | + | + | + |
| From ACC to left SMG | Increased | Increased | | + | | + |
| From ACC to right MFG | | Increased | Increased | + | - | - |
| From left SMG to PCC | Increased | Increased | | + | | + |
| From left SMG to left AG | Increased | Increased | | - | | + |
| From left SMG to right AG | Increased | | | - | + | + |
| **From left SMG to left MFG** | **Increased** | | **Decreased** | | **+** | **+** |
| From left SMG to right MFG | Increased | Increased | | - | + | + |
| From right SMG to PCC | | Increased | Increased | + | | - |

| | | | | | | |
|---|---|---|---|---|---|---|
| From left MFG to right AG | Increased | Increased | | + | + | + |
| From right MFG to left AI | Increased | Increased | | + | + | + |
| From right MFG to ACC | Increased | Increased | | - | + | |
| *Positive impact of estradiol is reduced in the presence of high progesterone levels:* | | | | | | |
| From left AG to left AI | Increased | Increased | | | + | - |
| From left AG to ACC | Increased | Increased | | + | + | - |
| From left MFG to right AI | Increased | Increased | | + | | - |
| From right MFG to right AI | Increased | | Decreased | + | + | - |
| From right MFG to left SMG | Increased | Increased | Decreased | + | | - |
| *Positive impact of higher estradiol levels is reversed by progesterone negative impact and/or combinatory effect:* | | | | | | |
| From PCC to right AG | | | Decreased | + | - | + |
| From PCC to left AI | | Decreased | | + | | + |
| From PCC to right SMG | Increased | | Decreased | + | | |
| From left AI to left AG | Increased | | | + | - | + |
| From left AI to right AG | Increased | | Decreased | + | - | - |
| From left AI to ACC | Increased | Increased | | + | - | |
| From left AI to right MFG | Increased | | Decreased | | - | - |
| From right AI to mPFC | | Increased | | | - | - |
| From ACC to right AI | Increased | | Decreased | | | - |
| From ACC to right SMG | Decreased | | | + | - | - |
| From left MFG to left AI | Increased | Increased | | + | - | + |
| From left MFG to ACC | | Decreased | Decreased | + | - | + |
| From left MFG to left SMG | | Decreased | Decreased | + | - | - |
| From right MFG to right AG | | Decreased | | + | | + |
| From right MFG to mPFC | | Decreased | Decreased | + | | - |

**Table 1. Summary of connections that showed cycle phases differences organized by hormonal relations.** Columns 2-4 correspond to cycle phase differences that survived a 75% posterior probability threshold. The hormonal modulation of these connections are detailed in columns 5-7, indicated by a + if relationship to the connectivity parameter was positive, and - if the relationship was negative. Connections that were able to predict cycle phase are marked in bold. E: estradiol, P: progesterone, E*P: estradiol and progesterone interaction.
